# Optimising a self-assembling peptide hydrogel as a Matrigel alternative for 3-dimensional mammary epithelial cell culture

**DOI:** 10.1101/2023.09.15.557894

**Authors:** Eliana Lingard, Siyuan Dong, Anna Hoyle, Ellen Appleton, Alis Hales, Eldhose Skaria, Craig Lawless, Simon Saadati, Aline F. Miller, Marco Domingos, Alberto Saiani, Joe Swift, Andrew P. Gilmore

## Abstract

Three-dimensional (3D) organoid models have been instrumental in understanding molecular mechanisms responsible for many cellular processes and diseases. However, established organic biomaterial scaffolds used for 3D hydrogel cultures, such as Matrigel, are biochemically complex and display significant batch variability, limiting reproducibility in experiments. Recently, there has been significant progress in the development of synthetic hydrogels for *in vitro* cell culture that are reproducible, mechanically tuneable, and biocompatible. Self-assembling peptide hydrogels (SAPHs) are synthetic biomaterials that can be engineered to be compatible with 3D cell culture. Here we investigate the ability of PeptiGel® SAPHs to model the mammary epithelial cell (MEC) microenvironment *in vitro*. The positively charged PeptiGel®Alpha4 supported MEC viability, but did not promote formation of polarised acini. Modifying the stiffness of PeptiGel® Alpha4 stimulated changes in MEC viability and changes in protein expression associated with altered MEC function, but did not fully recapitulate the morphologies of MECs grown in Matrigel. To supply the appropriate biochemical signals for MEC organoids, we supplemented PeptiGels® with laminin. Laminin was found to require negatively charged PeptiGel® Alpha7 for functionality, but was then able to provide appropriate signals for correct MEC polarisation and expression of characteristic proteins. Thus, optimisation of SAPH composition and mechanics allows tuning to support tissue-specific organoids.

## INTRODUCTION

The mammary gland undergoes dramatic changes during puberty, the menstrual cycle, pregnancy, lactation and involution. These changes involve complex molecular remodelling involving cells and their extracellular matrix (ECM), dysregulation of which is implicated in breast cancer [1, 2]. Many of the molecular networks regulating mammary gland development, remodelling and cancer are poorly defined. One important approach to study these processes is through *in vitro* organoid models, which combine some of the strengths of two-dimensional (2D) culture systems with *in vivo* systems [3]. An organoid model needs to mimic the key features of the tissue in question, although there are often limitations in how well this can be achieved [4]. Consequently, there is a significant need to develop materials that better recapitulate the biochemical and biomechanical properties of a tissue while being experimentally reproducible. It is also valuable to be able to independently manipulate the properties of a synthetic tissue in order to understand their function. The development of novel biomaterials has been a key to the improvement of tissue-specific models [5].

Mammary gland organoid models have largely relied on a reconstituted basement membrane (BM) extract, Matrigel [6]. First described in 1977 as a rich source of BM proteins [7], Matrigel was subsequently used as a hydrogel for three-dimensional (3D) mammary epithelial cell (MEC) cultures, promoting them to form polarised acini that recapitulated aspects of *in vivo* function [8]. Matrigel is derived from the Engelbreth-Holm-Swarm (EHS) mouse sarcoma and is enriched in ECM components like collagen-IV, laminin, entactin and perlecan [9]. These allow Matrigel to support MEC differentiation into organoids that produce milk proteins in response to prolactin [10]. Matrigel also contains numerous growth factors and cytokines which contribute to the behaviour of embedded cells [11]. This broad functionality and bioactivity mean it remains popular for 3D organoid cultures.

Matrigel and similar tumour-derived materials suffer from limitations as an organoid scaffold [12]. Matrigel is biochemically complex, containing approximately 1800 proteins identifiable by mass spectrometry [13, 14]. These include various growth factors and cytokines capable of affecting cell behaviour [15]. This complexity is exacerbated by variations between different production batches, both in total protein concentration and specific composition, resulting in inconsistency across experiments [9]. Matrigel contains tumorigenic factors that may stimulate pro-oncogenic cell behaviour, a potential issue for modelling non-tumorigenic tissue environments [16]. Matrigel’s mechanical properties also suffer from batch-to-batch variation, affecting cell behaviour independently from its composition [17, 18]. Matrigel is mechanically weak, and its mechanical properties are difficult to modify due to the interplay between matrix density and pore size with matrix stiffness. Finally, Matrigel is xenogenic and unrepresentative of human tissues, which introduces the risk of exposing cells to contaminants such as the immunogenic lactate dehydrogenase-elevating virus (LDV) [19].

The limitations of Matrigel have prompted development of cell scaffolds that offer more consistency, definition and tuneability. Synthetic polymers such as polyethylene-glycol (PEG) or polyacrylamide (PAM) are widely used. Their simple composition and flexible cross-linking chemistry make them amenable to specific cell culture applications [20]. Monomeric acrylamide is cytotoxic, making it incompatible for embedding cells for 3D culture, but PEG hydrogels have been successfully functionalised with the ECM mimetic peptide motifs for 3D MEC culture [21].

Self-assembling peptide hydrogels (SAPHs) are a class of promising synthetic scaffolds for 3D culture. SAPHs are short, synthetic, amphipathic peptides that self-assemble into fibrillar structures. These fibres entangle with each other in water to form hydrated, fibrillar scaffolds[8]. SAPHs are simple, well defined and reproducible, making them amenable to a variety of modifications [21, 22]. Furthermore, being peptide-based, SAPHs are inherently biocompatible [23]. Different peptide sequences allow for variation in hydrogel properties such as charge and stiffness, allowing properties to be selected for specific applications [9]. Here we investigate the potential for beta-sheet forming SAPH PeptiGels® to model MEC function *in vitro*. Both positively charged PeptiGel® Alpha4 (“Alpha4”) and negatively charged PeptiGel® Alpha7 (“Alpha7”) support MEC viability and proliferation. However, both Alpha4 and Alpha7 have limited bioactivity and were unable to promote complex MEC acinar morphogenesis. We investigated whether modifying SAPH properties through altering their mechanical stiffness and functionalising them with laminin-111 to form a hybrid hydrogel promoted MEC function. We found that the combination of laminin and negatively charged Alpha7 enabled MEC-specific protein expression and acinar morphology.

## RESULTS

### Alpha4 supports MEC viability but not acinar polarisation of MCF10A cells

Alpha4 is a positively charged SAPH that has previously been shown to maintain viable cultures of mouse MECs for up to 7 days [24]. It has also been shown to support cultures of human mesenchymal stem cells and rabbit synoviocytes [22, 25]. We therefore first assessed the ability of Alpha4 as a cell culture scaffold for the human, non-malignant mammary epithelial cell line MCF10A. MCF10A cells are a well-established model for normal mammary development and polarity, and their behaviour within 3D Matrigel is well characterised [26]. We encapsulated MCF10A cells in either Alpha4 or Matrigel, and allowed them to grow over 21 days, during which time we monitored viability, growth and morphology. Cells were viable in Alpha4 for at least 21 days, as shown by brightfield microscopy, and formed clusters that superficially resembled the 3D acinar structures formed in Matrigel (**Fig.1A**). We compared clusters grown in Alpha4 and Matrigel and measured their number, size and circularity (**Fig.1B**). 3D cultures were stained with DAPI at days 7, 14 and 21, and fluorescent images taken for image analysis. The number of acinar structures observed in Matrigel declined over 21 days, with 987 ±187 (+/- SEM) organoids counted at day 7 compared to 397 ± 255 organoids counted in Matrigel on day 21 (**Fig. 1B**). In contrast, the number of cell clusters in Alpha4 increased over 21 days from 96 ± 55 (+/- SEM) clusters at day 7 to 291 ± 267 clusters at day 21. We compared the morphology of organoids grown in Alpha4 and Matrigel at days 7, 14 and 21 by measuring their area and circularity. In Matrigel, MCF10A acini undergo growth arrest after around 14 days [27], with measurable cross sectional areas between 1000 and 9000 µm^2^ (**Fig.1B**). However, cell clusters in Alpha4 continued to grow, and were significantly larger than those in Matrigel at day 21. Clusters in Alpha4 also had significantly lower circularity compared to Matrigel, suggesting poorer acinar organisation (**Fig.1B**).

**Figure 1.**
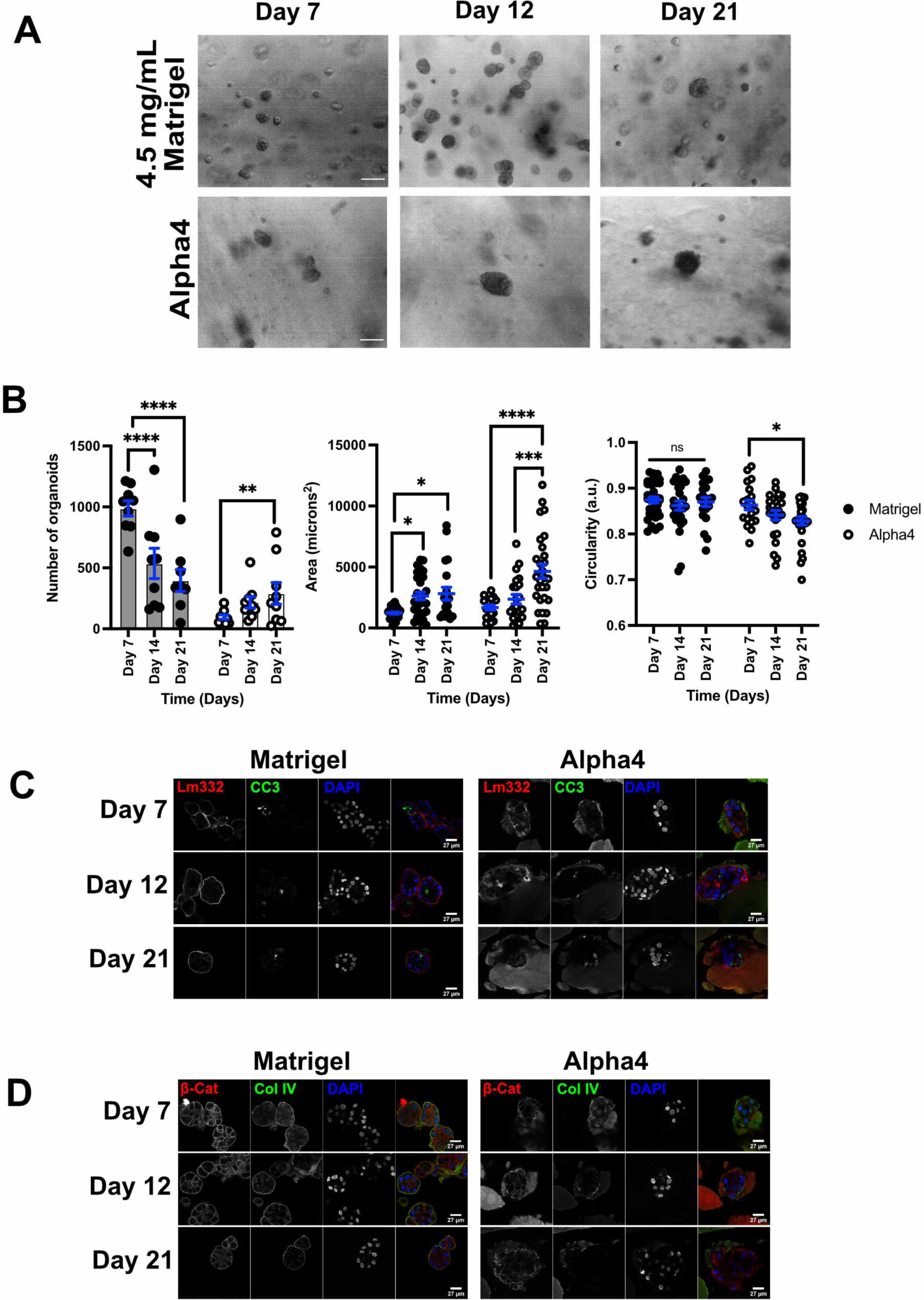
PeptiGel® Alpha4 alone supports MCF10A cell viability but does not allow 3D acinar organisation. A. Brightfield images of MCF10A cells encapsulated in Matrigel and Alpha4 (scale bars, 84 µm). MCF10A MECs were encapsulated into Matrigel or PeptiGel® Alpha4 for 7, 12 or 21 days. B. Comparison of organoid number, size and circularity between MCF10A cells embedded in Matrigel or Alpha4 for 7, 14 and 21 days. Experiments were repeated at least 3 times and data are shown as mean ± SEM (* P-value <0.05, ** p-value <0.01, *** p-value <0.001, **** p-value <0.0001) C. MCF10A cell organoids were grown in either Matrigel or Alpha4 for the indicated number of days. Organoids were extracted from the 3D cultures and immunostained for laminin-332 (Lm332) and cleaved caspase-3 (CC3). Nuclei were stained with DAPI (scale bars, 27 µm). D. MCF10A organoids as in C were immunostained for β-catenin (β-Cat) and collagen-IV (Col IV). Nuclei were stained with DAPI (scale bars, 27 µm).

To compare organisation of the 3D cell clusters in Matrigel and Alpha4, we examined localisation and expression of key markers of MCF10A organoid formation. Intact cell clusters were extracted from Matrigel or Alpha4 at days 7, 12 and 21, and fixed in paraformaldehyde. These were immunostained for laminin-332, collagen-IV, the cell-cell junction marker β-catenin, and cleaved caspase-3, a marker of apoptosis that occurs in the inner cells of MCF10A acini in Matrigel (**Fig.1C** and **D**). In Matrigel, MCF10A cells had a clearly defined BM surrounding the acini, containing laminin and collagen-IV. Cleaved caspase-3 was seen in cells within the intact acini, and β-catenin indicated cell-cell adherens junctions. In contrast, cell clusters from Alpha4 lacked an organised collagen-IV and laminin-332 BM, despite both proteins appearing to be made and secreted into the surroundings. Caspase-3 activity was broadly distributed, and b-catenin was seen, indicating cell-cell adherens junctions. Overall, these data revealed that although Alpha4 supports MCF10A viability, it is unable to fully recapitulate the MCF10A organisation seen in Matrigel.

### The mechanical properties of Alpha4 hydrogels can be tuned by dilution

ECM stiffness has been shown to affect a range of cellular behaviours, including regulation of gene expression and differentiation [6, 28]. We therefore asked whether the inability of Alpha4 to fully recapitulate MCF10A behaviour in Matrigel was due to differences in the stiffness of the two materials. We performed oscillatory shear rheometry on SAPH Alpha4 and 4.5 mg/mL Matrigel, using amplitude sweep analysis on three independent preparations of each gel. Gels were preconditioned in culture media at 37 °C and 5% CO_2_. Each preparation was measured three times at 1 Hz frequency within the linear viscoelastic region in the strain range 0.01 to 20% (**Fig. 2A**). Alpha4 showed a storage modulus (G’) of ∼ 400 Pa, although other data has also reported it to be in the region of 650 Pa [29]. In marked contrast, 4.5 mg/mL Matrigel had a G’ of ∼ 6 Pa. Published rheology data on Matrigel has reported widely different values of G’ that can be up to 5-fold stiffer, possibly reflecting batch variation [30]. However, Matrigel is considerably softer than Alpha4.

**Figure 2.**
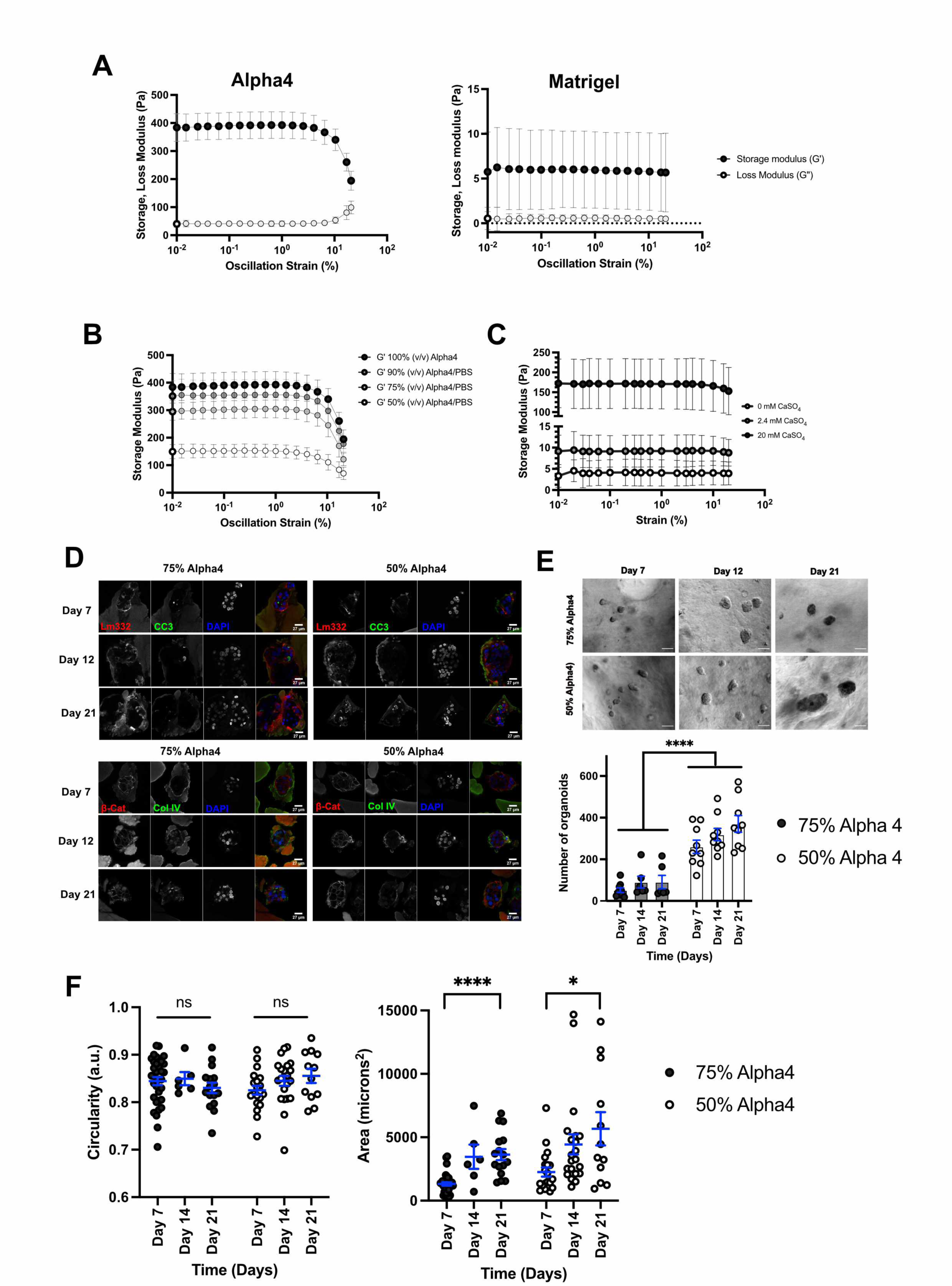
Modifying Alpha4 hydrogels stiffness increases organoid viability but does not alter their polarity. A. Oscillatory rheometry comparison of storage (G’) and loss modulus (G”) between Alpha4 and 4.5 mg/mL Matrigel. Measurements were taken at 1 Hz. B. Storage moduli at 1 Hz of PeptiGel® Alpha4 diluted with different volumes of PBS to the indicated proportions. All measurements were performed at least 3 times. Means ±SD are shown. C. Storage moduli at 1 Hz of Matrigel/alginate hydrogels stiffened using the indicated concentrations of calcium sulphate (CaSO_4_). Means ± SD are shown. All measurements were performed at least 3 times, but underloading of some Matrigel-alginate samples at 20 mM CaSO_4_ led to their exclusion from analysis. D. MCF10A cells were cultured within Alpha4 hydrogels at either 75% or 50% for 7, 12 or 21 days before organoids were isolated and immunostained for laminin-332 (Lm332), cleaved caspase-3 (CC3), β-catenin (b-Cat) and collagen-IV (Col IV), as indicated. Nuclei were stained with DAPI (scale bars, 27 µm). E. MCF10A organoid viability assessed in diluted PeptiGel®. Brightfield images of MCF10A cells encapsulated in 50% and 75% Alpha4 gels over 21 days (scale bars, 84 µm), and the number of organoids quantified at days 7, 14 and 21. Data are shown as mean ± SD. **** indicates P-value <0.0001 F. Organoid size and circularity of MCF10A cells cultured in 75% or 50% Alpha4 hydrogels for 7, 12 and 21 days. All measurements were performed at least 3 times. Data are shown as mean ± SD. P- values: * <0.05; ** <0.01; *** <0.001; **** <0.0001.

We considered the possibility that the stiffness of Alpha4 prevented MCF10A differentiation. Several studies have modified the stiffness of Matrigel to better mimic tissue mechanics. One way to achieve this is with an interpenetrating network of Matrigel and the polysaccharide alginate, which can be crosslinked with calcium ions to stiffen the network [31]. Matrigel-alginate gels with 2.4 mM and 20 mM CaSO_4_ have reported G’ values of around 80 and 310 Pa respectively, compared with 30 Pa without crosslinking [30]. Since Alpha4 is a porous network of peptides and water, we hypothesised that we could reduce the G’ by diluting it with phosphate-buffered saline (PBS). We found that Alpha4 could be diluted by up to 50% before the hydrogel began to fragment. This dilution limit was probably due to the peptide network unable to accommodate additional fluid. Amplitude sweeps showed that diluting Alpha4 with up to 50% (v/v) PBS reduced G’ to ∼ 150 Pa (**Fig. 2B**). Thus, Alpha4 can be reproducibly modified by dilution to create softer gels.

We compared these values for Alpha4 with Matrigel/alginate hydrogels. We performed rheology on soft (0 mM CaSO_4_), medium (2.4 mM CaSO_4_), and stiff (20 mM CaSO_4_) Matrigel/alginate gels at 1 Hz frequency (**Fig. 2C**). Matrigel/alginate hydrogels displayed elastic behaviour across the strain range 0.01-20% and increased in stiffness with CaSO_4_ addition. Soft Matrigel-alginate gels had a mean value of G’ of 4.2 Pa, similar to Matrigel alone. 2.4 mM CaSO_4_ increased mean G’ to 9.2 Pa. The stiffest Matrigel/alginate gels, with 20 mM CaSO_4_, showed a mean G’ of 171.8 Pa. As with Matrigel alone, these values are almost 5-fold lower than published measurements with similar Matrigel/alginate hydrogels. This data suggest that making comparisons between independent studies is difficult. Diluting the SAPH Alpha4 (with PBS) brought its G’ value into the range achievable with Matrigel.

### Softening the Alpha4 hydrogel by dilution is insufficient to cause acinar polarisation of MCF10A cells

We next examined the behaviour of MCF10A cells encapsulated in soft (50% v/v) and medium (75% v/v) Alpha4 hydrogels. Single MCF10A cells were encapsulated in each gel and grown for 7, 12 and 21 days, after which cell clusters were isolated, fixed and immunostained as before. Diluting Alpha4 did not improve the organisation of MCF10A acini. Localisation of laminin and collagen-IV both showed a similar distribution to that observed with cells grown in undiluted Alpha4. Although both ECM proteins did appear to be secreted, they were not organised into the defined BM seen with cells cultured in Matrigel (cf. **Fig.1C** and **D**). β-catenin and cleaved caspase-3 also showed a distribution similar to that in undiluted Alpha4. We quantified the number, size, and circularity of MCF10A clusters in 75% and 50% Alpha4 hydrogels. Reducing the stiffness of Alpha4 appeared to encourage increased cluster viability, in both the medium and soft gels compared to stiff Alpha4 (cf. **Fig.2E** and **Fig.1B**). Although we saw no differences in the circularity of cell clusters between any of the Alpha4 hydrogel preparations (**Fig.2F**), they did grow into larger organoids when the SAPH was softer.

To gain a holistic view of how softening the SAPH altered MCF10A cell function, and how this compared with cells cultured in Matrigel, we performed mass spectrometry (MS) analysis comparing acini grown in 75% and 50% Alpha4 with cells grown in 4.5 mg/mL Matrigel. MCF10A cells were encapsulated into each hydrogel and grown for 14 days, after which clusters were extracted and processed for liquid chromatography coupled tandem MS (LC-MS/MS). To exclude exogenous proteins originating from the media or hydrogel from the downstream comparison, we also analysed samples of cell-free Alpha4 and Matrigel. Approximately 800 – 1000 unique proteins were identified in each sample. Principal component analysis (**Fig.3A**) indicated clear separation between all three conditions, with the most separation observed between cells in Matrigel and those in either 75% or 50% Alpha4. However, there was separation between the two Alpha4 conditions, indicating that altering the mechanical stiffness of the SAPH altered MCF10A cell behaviour.

**Figure 3.**
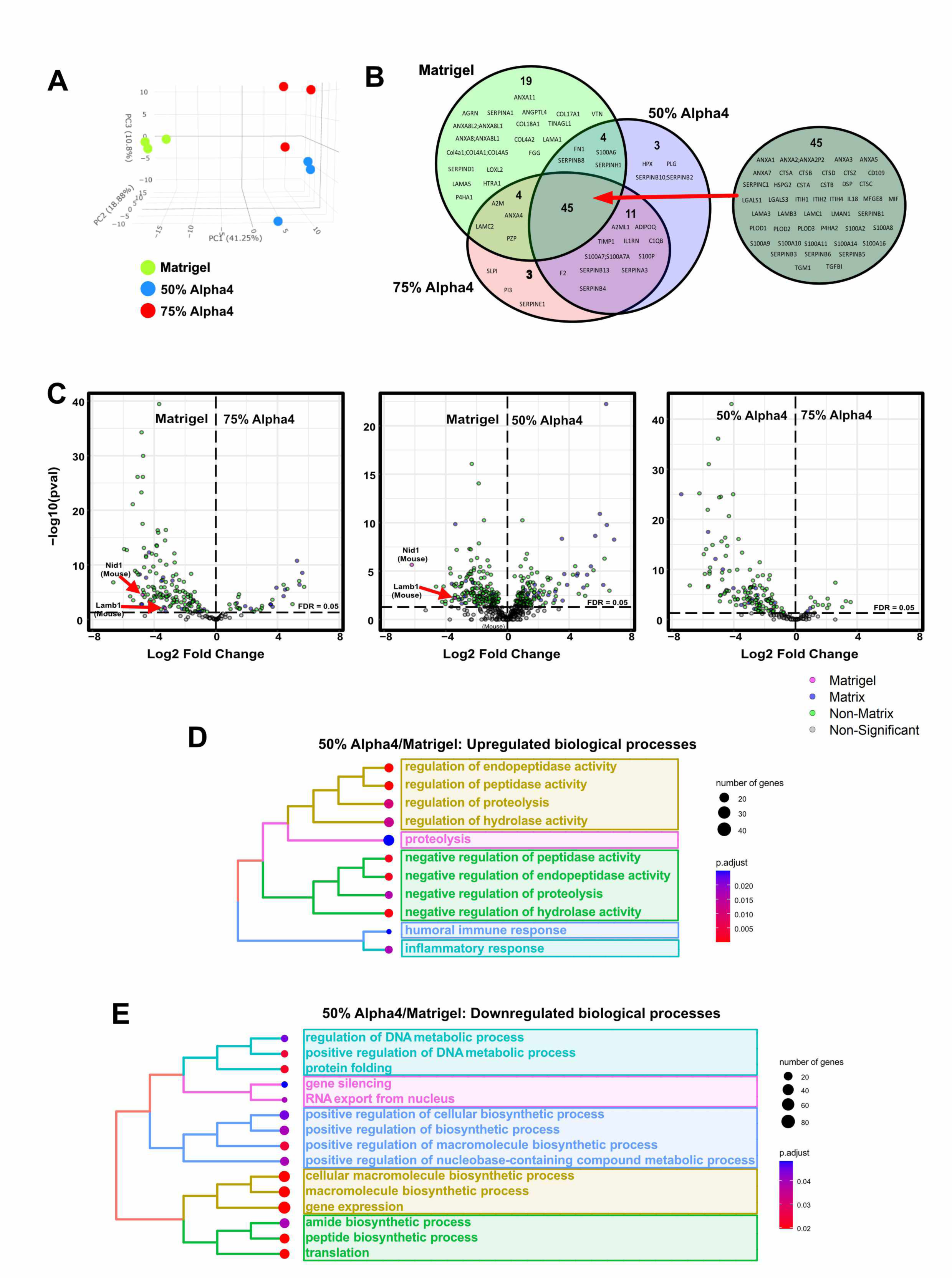
Mass spectrometry analysis of MCF10A cells encapsulated in Matrigel, stiff and soft Alpha4 identifies distinct proteomic profiles. A. Principal component analysis of proteomic analysis from of proteins quantified in cell-laden Matrigel, soft (50%) and medium (75%) Alpha4 gels. Three replicates from each condition were analysed. B. Venn diagram of mass spectrometry analysis showing unique and overlapping Matrisome proteins from each condition. C. Volcano scatter plot showing mass spectrometry comparison of proteins isolated from MCF10A organoids cultured in Matrigel, 75% Alpha4 and 50% Alpha4 hydrogels. Statistical significance (- log10 p-value) is shown versus magnitude of change (Log2 Fold Change). Plots depict upregulated (positive ratio) and downregulated (negative ratio) proteins. A positive Log2 change indicates proteins enriched in the condition named in the right hand panel of each plot. P-value calculated via MSqRob from three independent replicates (p <0.05). Matrigel specific proteins (mouse) are indicated, along with matrisome (Matrix) and other (Non-Matrix) proteins. D. Gene set enrichment analysis (GSEA) comparing MCF10A organoids grown in Matrigel with those grown in soft (50%) Alpha4, filtered for GO: biological processes. Indicates pathways that are upregulated in MCF10A cells encapsulated in 50% Alpha 4 relative to Matrigel. E. GSEA as in D but identifying pathways that are downregulated in MCF10A cells encapsulated in Alpha4 relative to Matrigel.

MCF10A cells grown in Matrigel make and organise a BM of laminin-5 and collagen-IV, whereas those grown in Alpha4 did not (**Fig.1C** and **D**). To further interrogate the ability of MCF10A cells to make ECM, we examined the LC-MS/MS data specifically for components of the Matrisome expressed under each condition (**Fig.3B**) [32]. 45 Matrisome components were identified in all three hydrogels, including the chains of laminin-5 (LAMA3, LAMB3), consistent with immunofluorescence (IF) data (c.f. **Fig.1C**, **D**). However, MCF10A cells in Matrigel expressed a more diverse set of ECM proteins, with 19 proteins not identified in either Alpha4 dilution. Cells in Matrigel made collagen-IV chains, COL4A1 and COL4A2, required for correct BM organisation and assembly, as well as COL4A5, collagens 17 and 18, and three other laminin isoforms. These data indicate that Matrigel provides a more permissive environment for MCF10A cells to differentiate and organise an appropriate ECM than does Alpha4.

To determine the effect of softening Alpha4 on MCF10A cell function we undertook quantitative comparison of the LC-MS/MS by analysing the mean fold-change and P-value for differentially identified proteins, examining total proteins and not just components of the Matrisome (**Fig.3C**). Comparison between cells within stiff Alpha4 with those in Matrigel showed that more proteins were significantly downregulated in the SAPH than were upregulated. In contrast, similar numbers of proteins were significantly upregulated and downregulated in soft Alpha4 gels in comparison with Matrigel. When comparing the two Alpha4 hydrogel cultures, there were significantly more downregulated proteins in the stiffer culture conditions. Thus, MCF10A cells are able to sense and respond to the mechanical properties of Alpha4 by regulating protein expression. Overall, these differences in protein expression between soft and stiff Alpha4 samples and Matrigel indicate that matrix stiffness and composition both contribute to MCF10A function.

Although Alpha4 diluted to 50% appeared to be the most favourable SAPH condition for MCF10A function, the protein expression profile was still distinct from cells in Matrigel. To understand the functional significance of these differences, we undertook gene ontology over representation analysis on the LC-MS/MS data to determine what key biological processes were different between cells grown in 50% Alpha4 and Matrigel. Using clusterProfiler’s gene ontology overrepresentation test, we identified significantly upregulated processes (p <0.05) enriched in soft Alpha4 gels in comparison with Matrigel (**Fig.3D**). Biological processes found to be significantly enriched in soft Alpha4 included those involved in regulating catabolic enzyme activity. Proteins associated with the humoral immune and inflammatory responses were detected, although these did not constitute the most enriched biological processes upregulated in soft Alpha4 samples. Downregulated biological processes mostly appeared to be affiliated with metabolic and housekeeping processes. These findings suggested that key difference in MCF10A cell behaviour in soft Alpha4 compared to Matrigel is a reduction in metabolic activity.

These data suggest that reducing the stiffness of Alpha4 improves viability and growth of MEC clusters, but that the mechanics alone are not sufficient to recapitulate differentiation and organisation of cells in Matrigel.

### MCF10A cells require laminin within the hydrogel scaffold to maintain acinar organisation

It was clear from the preceding data that MCF10A cells required specific biochemical signals from components of Matrigel that were not provided by a synthetic SAPH. Previous studies have shown that MECs form polarised acini within laminin containing hydrogels [33]. Furthermore, these allowed primary MECs to produce milk proteins in response to the lactogenic hormone, prolactin. We therefore asked whether MCF10A acini would maintain their organisation if embedded into either laminin or collagen gels. To initially test the response to collagen and laminin we grew MCF10A cells within 4.5 mg/mL Matrigel for seven days so that they had formed multicellular acini with an established BM (c.f. **Fig.4A** and **Fig.1**). Live, intact acini were then extracted from the Matrigel and embedded within secondary hydrogels, either Matrigel, Alpha4 SAPH, 1.5 mg/mL collagen-I or 6.1 mg/mL laminin-111. The concentrations of collagen-I and laminin-111 were chosen as these were able to form mechanically robust hydrogels capable of supporting cell cultures for at least seven days. Whereas acini in Matrigel and Alpha4 maintained their gross structure, those in collagen-I rapidly lost any organisation and cells became dispersed. In contrast, MCF10A acini in laminin hydrogels maintained a clearly defined organisation.

**Figure 4.**
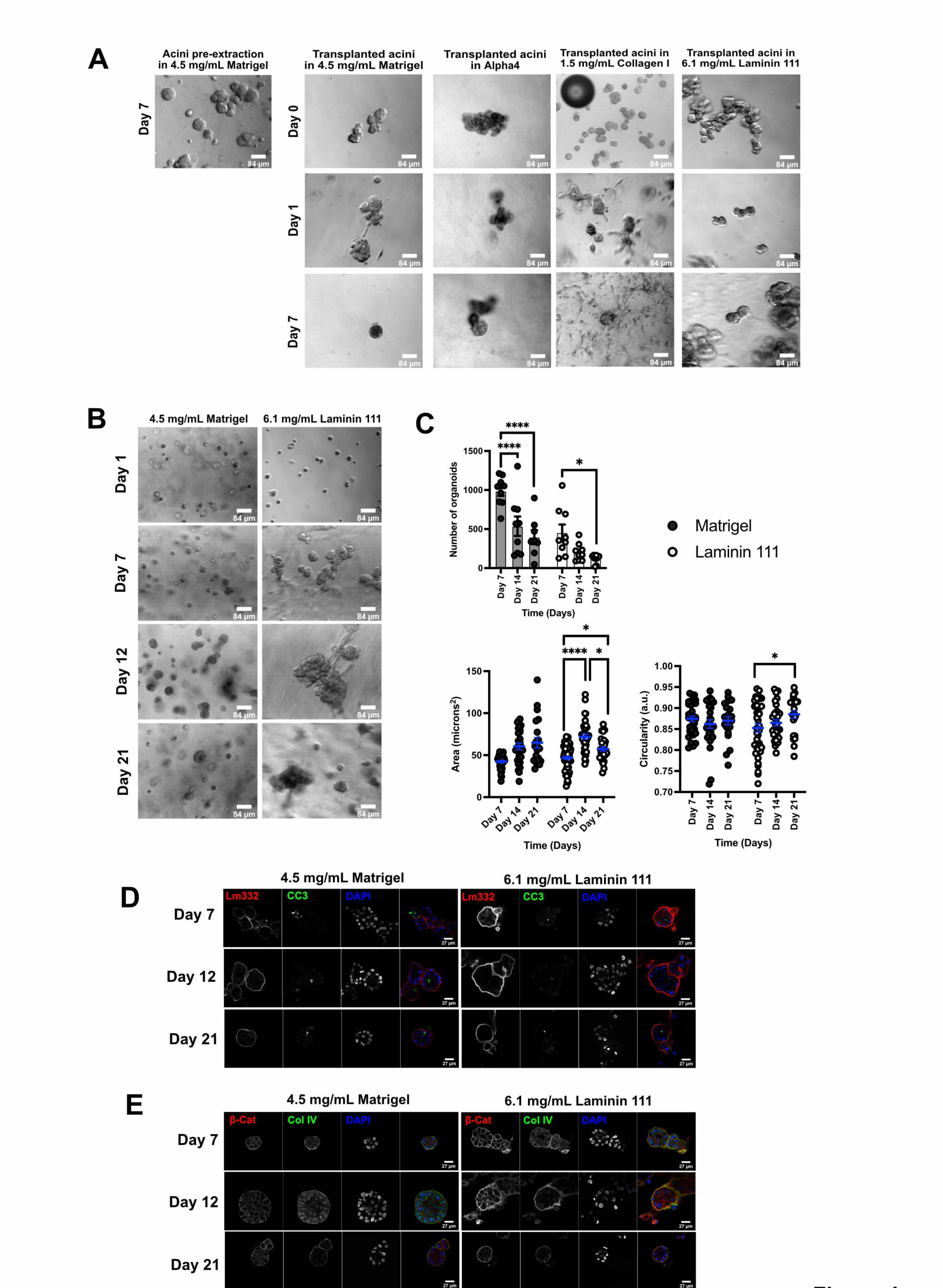
MCF10A acinar development and polarity requires laminin. A. MCF10A organoids initially grown in 4.5 mg/mL Matrigel for 7 days before being isolated and transplanted into either Matrigel, Alpha4, 1.5 mg/mL collagen-I or 6.1 mg/mL laminin-111. Brightfield images of the transplanted organoids at the indicated times. Scale bar = 84 µm. B. Brightfield images of single cell suspensions of MCF10A cells embedded in either 4.5 mg/mL Matrigel or 6.1 mg/mL laminin-111 and cultured for the indicated days. Scale bar = 84 µm. C. Quantification of organoid number, size and circularity of MCF10A cells grown in either Matrigel or laminin-111 for the indicated number of days. All measurements were performed at least 3 times. Data = mean ± SD. P-value: * <0.05; *** <0.001; **** <0.0001. D. MCF10A organoids grown in either Matrigel or laminin-111 for the indicated number of days. Organoids were extracted from the 3D cultures and immunostained for laminin-332 (Lm332) and cleaved caspase-3 (CC3). Nuclei were stained with DAPI (scale bars, 27 µm). E. MCF10A organoids as in D were immunostained for β-catenin (β-Cat) and collagen-IV (Col IV). Nuclei were stained with DAPI (scale bars, 27 µm).

To determine whether laminin-111 hydrogels were able to provide signals for acinar development, single MCF10A cells were encapsulated in laminin-111 gels and compared to cells in Matrigel. Cells were imaged over 21 days using brightfield and confocal IF imaging. Brightfield imaging showed acini forming in laminin-111 gels that were similar to those in Matrigel over the same time frame (**Fig.4B**). Some tubular structures were also observed in laminin-111 gels, which were not seen with cells grown in Matrigel. To measure organoids within the Matrigel and laminin, cultures were fixed and stained with DAPI on days 7, 14 and 21 days. Fluorescent images of organoids were then counted in QuPath (**Fig.4C**). Overall, fewer organoids were growing within the laminin gels compared to Matrigel. The total number of organoids decreased over the 21-day period in both gels. We quantified the size and circularity of the organoids. There was little difference in acini between the Matrigel and laminin cultures (**Fig.4C**). Acini grew in size in both Matrigel and laminin between days 7 and 14. There was no significant difference in circularity was found between Matrigel and laminin 111 cultures. Any minor differences might be explained by the more complex composition of Matrigel regarding growth factors, although the laminin-111 used was purified from Matrigel.

We encapsulated single MCF10A cells in either 4.5 mg/mL Matrigel or 6.1 mg/mL laminin-111 for 7, 12 and 21 days. Acini were then extracted, fixed and immunostained for laminin-332, cleaved caspase-3, β-catenin and collagen-IV (**Fig.4D** and **E**). Acini from laminin-111 hydrogels were indistinguishable from those in Matrigel and displayed a clearly defined BM, seen through both laminin-332 and collagen-IV localisation. β-catenin showed clearly defined cell-cell junctions, and cleaved caspase-3 positive cells indicated apoptosis occurring within the central lumen of the developing structures. The similarity of MCF10A acini in laminin-111 compared to Matrigel indicate that it alone can provide the appropriate biochemical and mechanical cues for MEC culture.

### Laminin functionalises SAPHs to support MCF10A acinar development

Laminin-111 provided an appropriate biochemical and biomechanical microenvironment to support MCF10A cells to organise into the same polarised, acinar structures as seen in Matrigel. However, purified laminin cannot be mechanically tuned and requires a high concentration to form a hydrogel, making it impractical as a 3D culture scaffold. We therefore asked if supplementing SAPHs such as Alpha4 with lower concentrations of laminin would combine both the required mechanics and a suitable biochemical signal. We therefore encapsulated MCF10A cells within either Matrigel, laminin- 111, Alpha4 or Alpha4 supplemented with 1 mg/mL laminin-111. We also included a comparison with Alpha4 diluted with Matrigel at a final concentration of 1.5 mg/mL. Cultures were maintained for seven days before imaging (**Fig.5A**). MCF10A cells formed organoids as previously seen within Matrigel, laminin-111 and Alpha4. Including 1.5 mg/mL Matrigel within Alpha4 gel promoted acinar development, although the structures appeared confined to discrete areas within the hydrogel, suggesting that Matrigel and Alpha4 did not fully mix. Notably, addition of laminin to Alpha4 did not stimulate acinar development in MCF10A cells. Furthermore, there was a precipitate within Alpha4 combined with laminin, suggesting that they were not fully compatible. A possible clue as to why Alpha4-laminin hydrogels contained a precipitate is that Alpha4 has an overall positive charge. We therefore asked if the soft, negatively charged SAPH, Alpha7, would be compatible with laminin. We encapsulated MCF10A cells for seven days within either Alpha7 alone, Alpha7 with Matrigel or Alpha7 with 1 mg/mL laminin (**Fig.5B**). MCF10a cells were viable in Alpha7 alone but appeared mostly to be single cells. In contrast, addition of either Matrigel or laminin increased the number of acinar-like structures formed over seven days.

**Fig. 5.**
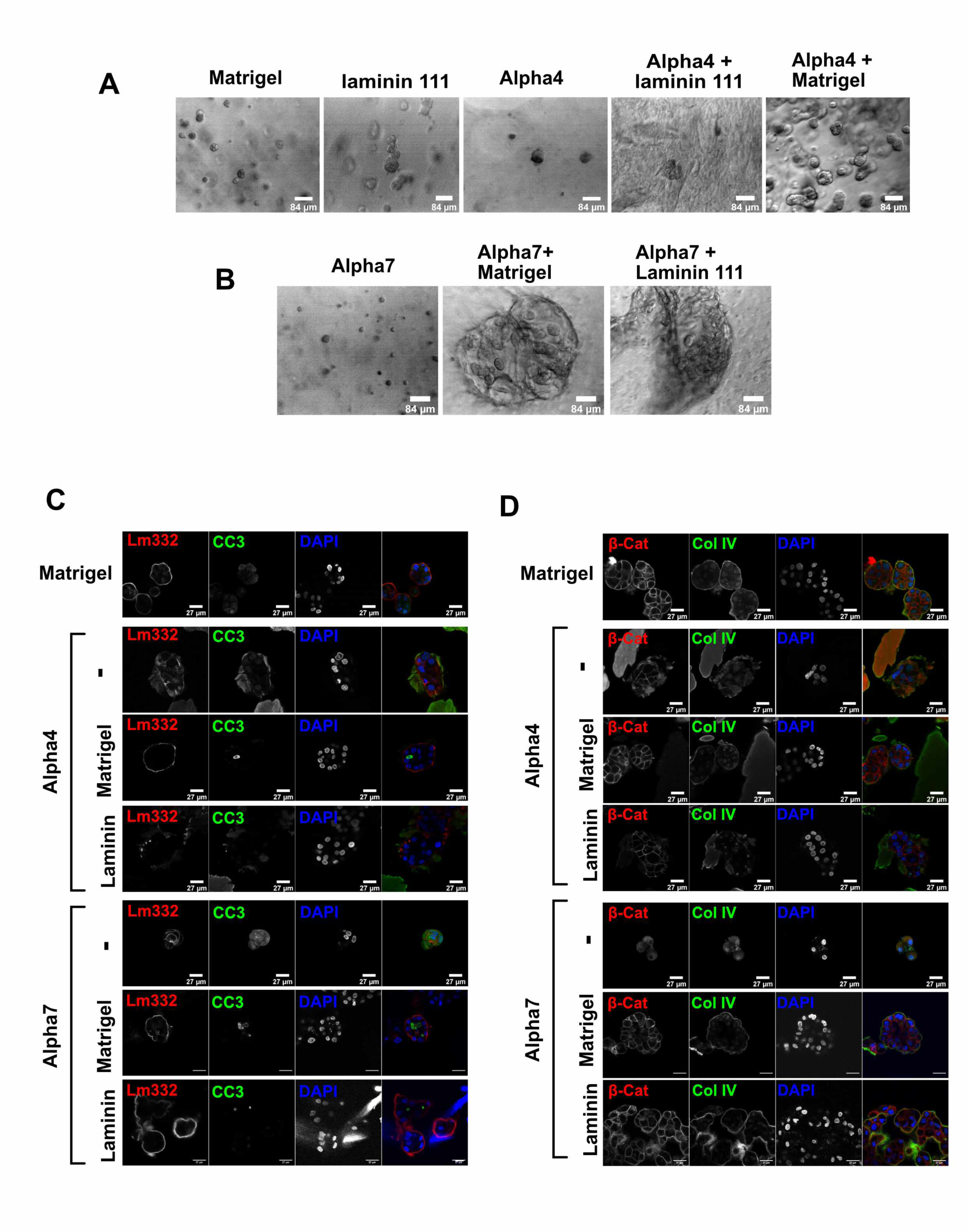
SAPHs functionalised with laminin support MCF10A cell acinar development. A. Comparison of MCF10A organoids grown for 7 days from single cell suspensions in Matrigel, laminin-111, Alpha4, or Alpha4 supplemented with either Matrigel or laminin-111. Brightfield images show lack of organoids in Alpha4 gels combined with laminin. Scale bar = 84 µm. B. MCF10A organoids grown from 7 days in Alpha7 alone or supplemented with Matrigel or laminin-111. Scale bar = 84 µm. C. MCF10A organoids grown in either Matrigel, Alpha4 or Alpha7. Alpha4 and Alpha 7 were supplemented with either laminin-111 or Matrigel as indicated. Organoids were cultured for 7 days, extracted from the 3D cultures and immunostained for laminin-V (Lm332) and cleaved caspase-3 (CC3). Nuclei were stained with DAPI. Scale bars = 27 µm. D. MCF10A organoids as in D were immunostained for β-catenin (b-Cat) and collagen-IV (Col IV). Nuclei were stained with DAPI. Scale bars = 27 µm.

We compared the functionalisation of both Alpha4 and Alpha7 with Matrigel or laminin by encapsulating single MCF10A cells into each hydrogel or composite and culturing them for seven days. Acini were then isolated, fixed, and immunostained for laminin, cleaved caspase-3, collagen- IV and β-catenin (**Fig.5C** and **D**). Compared with Matrigel alone, Alpha4 did not support correct organisation of MCF10A acini when supplemented with laminin, although supplementing with Matrigel did. Alpha7 alone was unable to support acinar organisation, and MCF10A cells were seen as very small cell clusters with no BM deposition. However, Alpha7 supplemented with laminin promoted acini that were indistinguishable from those in Matrigel. Indeed, these acini showed clear deposition of a BM with both laminin-332 and collagen-IV, identical to that of cells cultured in Matrigel.

To determine how laminin altered behaviour of MCF10A cells within Alpha7 we performed an unbiased proteomic analysis. Triplicate cultures of MCF10A cells were encapsulated within either Alpha7 or Alpha7 supplemented with 1 mg/mL laminin-111 for 14 days. To eliminate proteins within the culture media and the laminin-111 we also analysed cell-free preparations of each scaffold, with any proteins with the same abundance in both blank and cell-containing sample removed from downstream analysis. Over 2000 proteins were detected by LC-MS/MS from each sample from both the Alpha7 and Alpha7/laminin. After normalisation of peptide intensity, principal component analysis clearly separated lysates obtained from Alpha7 and Alpha7/laminin gels (**Fig.6A**). As expected from the dramatic difference in acinar organisation (**Fig.5C** and **D**), this clearly shows that supplementing laminin-111 to Alpha7 changes protein expression in MCF10a cells. Quantitative comparison of the LC-MS/MS data for significant changes in protein expression indicated broad differences in protein expression between the Alpha7 and Alpha7/laminin (**Fig.6B**). In order to identify differentially regulated biological processes regulated by laminin we used clusterProfiler’s gene ontology overrepresentation test. This identified numerous significantly upregulated processes (p <0.05) when laminin was added to Alpha7 (**Fig.6C**). In contrast to Alpha4 hydrogels, these included processes associated with epithelial cell differentiation, development and processes expected if MCF10A cells were displaying a phenotype indicative of MEC differentiation. Downregulated biological processes included DNA replication, mRNA processing, and mitosis. This is consistent with the smaller size and lack of BM organisation of cell clusters in Alpha7, overall suggesting that MCF10A cells are less proliferative in the absence of laminin (**Fig.6D**).

**Figure 6.**
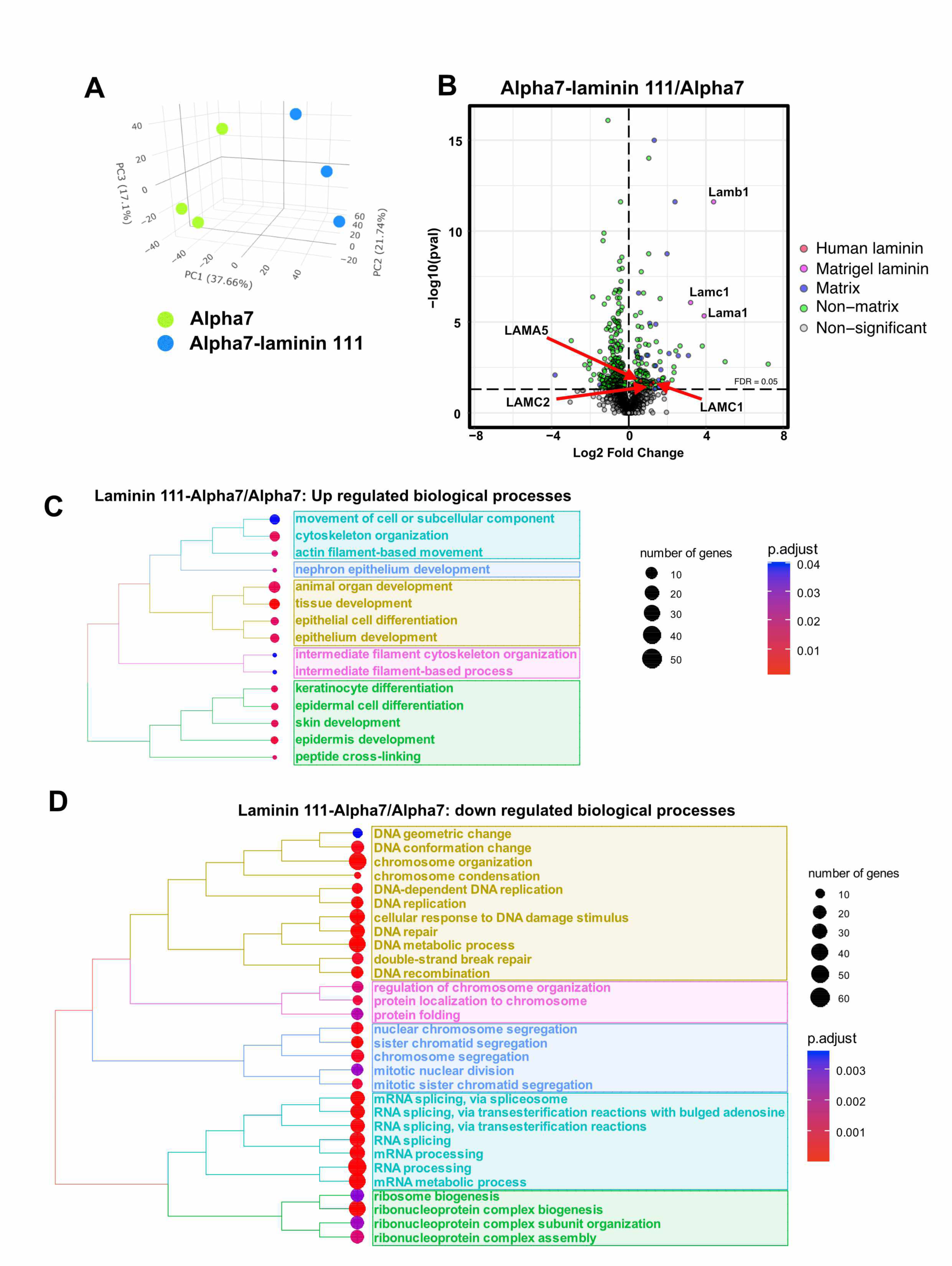
Mass spectrometry analysis of MCF10A organoids grown in Alpha7 with laminin show protein expression indicating MEC differentiation. MCF10A organoids were grown for 7 days in either Alpha7 hydrogels or Alpha7 supplemented with laminin-111. Proteins were isolated and quantified by mass spectrometry. Three independent replicates were analysed. A. Principal component analysis of organoids grown in Alpha7 and Alpha7/laminin show separation between the two conditions. B. Volcano scatter plot showing mass spectrometry comparison of proteins isolated from MCF10A organoids in Alpha7/laminin hydrogels versus Alpha7 hydrogels. Statistical significance (-log10 p- value) is shown versus magnitude of change (Log2 Fold Change). Plots depict upregulated (positive ratio) and downregulated (negative ratio) proteins. A positive Log2 change indicates protein abundance is enriched in the Alpha7-Laminin111 condition. P-values calculated using MSqRob from three independent replicates (p <0.05). Mouse laminin (Lama1, Lambb1 Lambc1, from Matrigel) are indicated. Human laminin isoforms identified (LAMA5, LAMC1, LAMC2) show increased abundance in Alpha7/laminin hydrogels versus Alpha7 hydrogels. C. Gene set enrichment analysis (GSEA) conducted on the data in B. showing GO:biological processes upregulated in MCF10A cells in Alpha7/laminin hydrogels versus Alpha7 hydrogels. Key process include various terms associated with epithelial differentiation. D. GSEA conducted on the data in B. showing GO:biological processes downregulated in MCF10A cells in Alpha7/laminin hydrogels versus Alpha7 hydrogels. Relationships between groups are displayed by connecting lines.

Together, these data show that optimisation of SAPHs as a mechanical scaffold along with appropriate supplemental microenvironmental components such as tissue specific ECM proteins allows for a reproducible alternative to Matrigel for MEC culture models.

## DISCUSSION

In this study we have demonstrated that SAPHs Alpha4 and Alpha7 support viability of mammary epithelial cells in 3D culture models. However, the properties of these hydrogels need optimisation to replace established culture conditions using organic matrices like Matrigel. SAPHs provide a biocompatible and chemically defined hydrogel scaffolds which are extremely consistent between batches. Different peptides, however, have widely different properties, including their charge and mechanical stiffness. The results presented here show that optimising SAPHs is important to fully utilise these in 3D tissue culture models.

Previous studies found neutral charge SAPH was compatible with MCF7 and MDA-MB-231 breast cancer models, recapitulating features such as tumour hypoxia and drug accessibility [34]. In contrast, neutral charged SAPHs did not support MCF10A cell viability or growth (data not shown), whereas charged peptides such as Alpha4 did. However, although MCF10A cells were viable and proliferated within Alpha4, they failed to recapitulate their polarisation seen in Matrigel. One aspect of SAPHs determined by their peptide sequence is mechanical stiffness. MECs and breast cancer cells have been shown to respond significantly to ECM stiffness [35, 36]. Matrigel is both significantly softer that SAPHs and inconsistent between batches. We found Matrigel to have G’ ∼ 6 Pa, and this could be stiffened using an interpenetrating alginate network to ∼ 170 Pa. Other studies that have used Matrigel/alginate networks have reported values for G’ between 30 and 310 Pa using the same CaSO_4_ concentrations [37]. Our measurements for Alpha4 are consistent with reported values. The stiffness of SAPHs can be tuned through the amphiphilicity of the sequence altering the interactions between fibres. Utilising different SAPH sequences demonstrated that pancreatic cancer cells showed reduced apoptosis within a stiff hydrogel (PeptiGel® Alpha2, G’ ∼10 kPa) compared to a softer one (PeptiGel® Gamma2, G’ ∼4 kPa) [38]. We were able to modify the stiffness of Alpha4 by diluting it in an aqueous buffer. This had the effect of reproducibly softening Alpha4, avoiding the necessity for different peptide sequences and corresponding changes in charge. The G’ of Alpha4 could be reduced approximately 2.5-fold before the network became unstable. Proteomic analysis demonstrated that MCF10A cells within Alpha4 hydrogels synthesised epithelial specific ECM components, including collagen IV and laminin. However, although softening Alpha4 did alter gene expression in MECs, this alone did not recapitulate the protein expression or polarisation seen with organoids grown in Matrigel. Overall, this suggests that key biochemical signals are required to form an organised BM.

Laminin-111, the major component of Matrigel, can promote primary MECs differentiation in 3D cultures [29]. High concentration purified laminin-111 formed hydrogels which promoted 3D acinar polarisation of MCF10A cells. We were able to combine laminin at lower concentrations within SAPHs, but this was then only compatible with negatively charged Alpha7 peptide. Laminin appeared to precipitate within Alpha4 and failed to support MEC acinar organisation. One possible explanation is that laminin compatibility is mediated by electrostatic interactions with the charged SAPH. Electrostatic interactions between cells and their environment regulate viability, differentiation, and adhesion. Positively charged SAPHs promote cell attachment via interactions between negative charges on the cell surface and positively charged scaffold [39–41]. Conversely, negatively charged hydrogels can repel negatively charged cell surface moieties, resulting in cells remaining single and rounded, as we observed in non-functionalised Alpha7 [42]. Electrostatic interactions between oppositely charged moieties are involved in laminin binding, where anionic glycans presented on dystroglycan [43, 44] form salt bridges with cationic lysine residues within the laminin globular domains [4, 45]. These interactions drive laminin polymerisation and BM formation [41, 42, 46]. Thus, the peptide sequence of SAPHs may be critical for how macromolecules interact within the hydrogel, and how they are presented to cells to provide key functionality. Consideration of the individual properties of SAPHs would appear to be essential for successful functionalisation.

A SAPH based on Alpha7 and functionalised with laminin recapitulated the MCF10A acinar organisation seen in complete Matrigel. Proteomic analysis indicated differentiation of MCF10A cells, with upregulation of BM components, as well as regulators of MEC differentiation such as Stat6 and interferon regulatory factor 6 [47, 48]. Pathway enrichment analysis highlighted several biological processes were upregulated that indicate mammary morphogenesis. Overall, these results show that SAPHs can be functionalised for MEC culture, creating a scaffold that is defined, simple and modifiable. Although here we used purified laminin, a fully synthetic functionalised hydrogel could be achieved by using recombinant full-length laminin or laminin fragments.

## MATERIALS AND METHODS

### Cell culture

Immortalised, non-tumorigenic human mammary epithelial cells (MCF10A) were sourced from ATCC and maintained in DMEM-F12 media supplemented with 5% filtered horse serum, 10 μg/mL insulin, 0.5 μg/mL hydrocortisone, 100 ng/mL cholera toxin and 20 ng/mL EGF. Cells were passaged at 70- 90% confluency.

PeptiGels® were purchased from Manchester BIOGEL (Alderley Park, UK) and are now sold by Cell Guidance Systems (Cambridge, UK). Matrigel and collagen I were purchased from Corning Life Sciences. High-concentration Cultrex 3-D Culture Matrix laminin-I was purchased from Trevigen.

For encapsulation of MCF10A cells into 3D gels, wells of a 24-well plate were coated with a 50 µL layer of either undiluted Matrigel (8.9 mg/mL) high-concentration laminin-I (6.1 mg/mL) and left to set for 30 minutes at 37°C. Passaged MCF10A cells were resuspended in 1 mL of media. For Matrigel, cell suspensions were mixed into blank DMEM to give a volume of 49.5 µL per gel, and 50.6 µL of 8.9 mg/mL Matrigel added to give a final total protein concentration of 4.5 mg/mL and cell density of 0.5 x 10^5^ cells per 100 µL of gel. 100 µL of the Matrigel-cell-DMEM solution was then added to each well before polymerising at 37°C/5% CO2 for 30 minutes. For laminin I gels, MCF10a cells in culture media were mixed with 6.1 mg/mL laminin-111 to a final concentration of 5.49 mg/mL and a cell density of 0.5 x 10^5^ cells per 100 µL of gel. 100 µL of laminin-cell mixture was then pipetted into each well and left to polymerise at 37°C (5% CO2) for 30 minutes. The gels were bathed in media which was refreshed every 2-4 days.

For collagen gels, collagen-I at a final concentration of 1.5 mg/mL (prepared from 3.98 mg/mL collagen I, 104 µL DMEM and 20 µL 10X Roswell Park Memorial Institute (RPMI) media, neutralised by adding 3 µL of 1M sodium hydroxide) was spread over the bottom surfaces of 24-well plates and left to set for 30 minutes at 37°C/5% CO_2_. MCF10A cells were prepared in 200 µL 1.5 mg/mL collagen-I prepared as above. The gels were incubated for 30 minutes and bathed in media which was refreshed every 2-4 days.

For SAPH cultures, 50 µL of PeptiGel® at room temperature were spread over the bottom surface of wells in 24-well plates. MCF10A cells were encapsulated via gentle pipetting into appropriate volumes of SAPH, with a final cell density of 0.5 x 10^5^ cells per mL. Following encapsulation, 100 μL of cell-laden hydrogels were pipetted into each well on top of the gel layer. Culture media was added and refreshed every 2-4 days.

To isolate intact organoids for either re-encapsulation in hydrogels or for immunofluorescent staining, 3D cultures were washed with 1 mL of PBS following removal of media and then depolymerised using 1 mL of ice-cold cell recovery solution (Corning) for 1 hour at 4°C. The depolymerised gels were resuspended, collected into tubes pre-coated with 1% BSA in PBS (w/v) and washed via centrifugation in PBS at 70 RCF. The supernatants were discarded, and the organoid pellets resuspended for re-encapsulation or fixed for staining.

### Oscillatory shear rheometry

The storage modulus of gels was measured using a Discovery HR-2 hybrid rheometer (TA Instruments, USA) with a 20 millimetre (mm) parallel plate and a gap size of 500 μM. Samples were prepared by aliquoting 180 μL of gel into ThinCert well inserts (1 μM pore size, Greiner Bio-One Ltd, Gloucestershire, UK). 900 μL of assay media was pipetted into the wells after 5 minutes and left to recover for 5 minutes before 100 μL of media was added to each insert. The gels were incubated at 37°C (5% CO_2_) for at least 30 minutes prior to testing. Following media exposure, samples were removed from the inserts by peeling off the bottom membrane of the insert and transferred onto the rheometer plate as described by Ligorio et al. [49]. The upper rheometer head was then lowered to the gap size and samples were equilibrated for 3 minutes at 37°C. Oscillatory amplitude experiments were performed at 1 Hz frequency and within the linear viscoelastic region in the strain range: 0.01 to 20%. The mean storage modulus values described in the results section were obtained at 0.2% oscillation strain. All measurements were repeated three times to ensure reproducibility.

### Immunofluorescent staining and microscopy

Extracted MCF10A organoids were fixed for 45 minutes in 4% formaldehyde in PBS (v/v) at room temperature, diluted with 10 mL of PBS and centrifuged at 70 RCF for 3 minutes at 4°C. The pellets were resuspended in 1 mL of organoid wash buffer (PBS containing 0.1% Triton-X-100 and 0.2% BSA), transferred to pre-coated, low adherent 24-well plates (Greiner Bio-One) and blocked for 15 minutes. Cell clusters were incubated with 2X primary antibodies in organoid wash buffer overnight at 4°C. The plates were retrieved, washed three times in organoid wash buffer for 1 hour at 4°C. Clusters were incubated with 2X secondary antibody solutions in organoid wash buffer overnight at 4°C, incubated with 2 μg/mL DAPI in PBS for 10 minutes on an orbital shaker before being washed 3 times with organoid wash buffers described above. Organoids were diluted in PBS and transferred to 6-well plates for imaging. Confocal images were collected on a Leica TCS SP8 AOBS upright confocal using a 63x/0.90 water immersion objective. The confocal settings were as follows, pinhole 1 airy unit, scan speed 400 Hz unidirectional, format 1024 x 1024. Images were collected using hybrid and photomultiplier detectors with the following detection mirror settings; DAPI 410-475 nm; Alexa-488 507-580 nm; Alexa-594 605-750 nm using the 405 nm (50%), 490 nm (30%) and 590 nm (30%) laser lines respectively. When it was not possible to eliminate crosstalk between channels, the images were collected sequentially. The acquired images were processed using ImageJ.

Primary antibodies used for immunofluorescent staining were: active caspase-3 (Rabbit, R&D Systems, AF835); laminin alpha3 chain (Mouse, R&D Systems, MAB21441); collagen IV (Rabbit,

Abcam, ab6586); β-catenin (Mouse, BD Biosciences, 610154). Secondary antibodies were from Invitrogen (donkey anti-mouse AlexaFluor 594, A21203; donkey anti-rabbit AlexaFluor 488 A21206).

### Assessment of MCF10A cell clusters

To assess organoid size and density, encapsulated MCF10A organoid hydrogels were prepared in triplicate as above and cultured for up to 21 days and fixed on either day 7, 14 or 21 as above. Fixed clusters were then stained with 1 μg/mL DAPI in PBS for 10 minutes, washed with 3D IF wash buffer for 10 minutes and then double-distilled water overnight. Fluorescent images of DAPI-stained clusters were collected as Z-stacks on an EVOS M7000 Imaging system (Thermo Fisher Scientific) using a 4X PlanFL 0.13 NA objective using the DAPI light cube. The acquired images were processed in QuPath 0.2.3 by manually counting fluorescent nuclei. Measurements were exported to GraphPad Prism for statistical analysis.

Clusters formed in Matrigel and laminin-111 were analysed in ImageJ to assess their area and circularity. Clusters in focus were traced around their periphery using the freehand tool (to measure circularity) or the freehand line tool (to measure area). The tracing was done by hand using a Wacom One drawing tablet and pen. Measurements were exported to GraphPad Prism.

### Mass spectrometry analysis

Hydrogel-encapsulated MCF10A cell organoids were lysed in 100 µL of 1X radioimmunoprecipitation (RIPA) buffer (50 mM Tris-HCL/pH 7.4, 150 mM sodium chloride, 1% IGEPAL, 0.1% (w/v) SDS, 1% sodium deoxycholate, 20 mM sodium fluoride, 2 mM sodium orthovanadate, 1X protease inhibitor cocktail). Following 15 minutes incubation on ice, the samples were sonicated for 180 seconds using a Covaris S220 ultrasonicator before being centrifuged at 3220 x g for 5 minutes at 4°C. Lysates were stored at 20°C prior to use.

Lysates were prepared for mass spectrometry using SDS-PAGE electrophoresis. Samples were allowed to migrate past the wells of a pre-cast 4-20% polyacrylamide gel. Once stained with InstantBlue, gel sections were submitted for peptide identification analysis to the University of Manchester Biological Mass Spectrometry Core Facility. The samples underwent in-gel digestion before being analysed using the Thermo Orbitrap Elite mass spectrometer coupled with a Thermo nanoRSLC system.

Raw data files were processed in MaxQuant (v 2.0.1.0) [50]. Spectra were searched against the Human (Homo Sapiens) reference proteome obtained from Uniprot (June 2021), which was modified to include the three murine laminin I subunit (LAMA1_MOUSE, LAMB1_MOUSE, LAMC1_MOUSE) peptide sequences obtained from the Mouse (Mus musculus) reference proteome (July 2021) [51]. Methionine oxidation and N-terminal acetylation were set as variable modifications and cysteine carbamidomethylation was set as a fixed modification. Fast label-free quantification was enabled, with a minimum label ratio of 2 selected. A minimum of 3 and an average of six sample neighbours were also set. Precursor tolerance for the first and main searches was set at 20 ppm and 4.5 ppm, respectively. MS/MS tolerance was set at 20 ppm, with a maximum of two missed cleavages allowed. The false discovery rate (FDR) of PSM and protein were set at 0.01 and “Match between runs” was enabled. MaxQuant peptides.txt and proteinGroups.txt output files were processed in R (v 4.1.2). Contrasts were set up and run using MsqRob2 (v 1.2.0) [52], where only proteins of FDR ≤0.05 were considered significant.

Gene ontology enrichment analysis was performed using the clusterProfiler package (v 4.0) [53]. Separate lists of significantly upregulated and downregulated ENTREZID gene IDs were queried against the org.Hs.eg.db (v 3.14.0) database. Overrepresentation analysis was performed and corrected for FDR using the Benjamani-Hochberg method by clusterProfiler to determine enriched genes in the lists. Only genes of p ≤0.05 were considered significant. Pathway enrichment analysis was then performed using ReactomePA (v1.38.0) [54], where separate lists of significantly upregulated and downregulated ENTREZID gene IDs were queried against the org.Hs.eg.db database and subject to overrepresentation analysis using the Benjamani-Hochberg method to determine enriched pathways and interaction networks. Only genes of p ≤0.05 were considered significant. All enriched genes and pathways were visualised in R using enrichplot (v 1.14.1).

### Statistical analysis

All quantitative values are presented as mean ± standard deviation. Data was analysed using one-way or two-way analysis of variance (ANOVA) in GraphPad Prism v9.3.0.

## ACKNOWLEDGEMENTS

The project was funded through the UK Regenerative Medicine Platform grant “Acellular/Smart Materials – 3D Architecture” (MR/R015651/1). EL was supported by a PhD scholarship from FBMH and Manchester BIOGEL. AH and EA were supported by studentships from the Biotechnology and Biological Sciences Research Council (BBSRC) and Wellcome Trust, respectively. JS was supported by a BBSRC David Phillips Fellowship (BB/L024551/1). ES and the Wellcome Trust Centre for Cell-Matrix Research were funded by the Wellcome Trust (088785/Z/09/Z). Proteomics was performed by the Biological Mass Spectrometry Core Facility (supported by BBSRC, Wellcome and the University of Manchester Strategic Fund). This work was supported in part by the International Alliance for Cancer Early Detection, an alliance between Cancer Research UK, Canary Center at Stanford University, the University of Cambridge, OHSU Knight Cancer Institute, University College London and the University of Manchester. Work in the Royce Institute is funded through EP/R00661X/1, EP/S019367/1, EP/P025021/1 and EP/P025498/1.

## AUTHOR CONTRIBUTIONS

Conceptualization, JS, AS, MD and APG; Investigation, EL, SD, AH, ES; Data Analysis, EL, EA, ES, SS, AH, CL; Writing – Original Draft, EL, APG; Writing – Visualization, Review & Editing, JS, MD, SS, AS, APG, EL; Project Administration and Funding Acquisition, APG, AS, JS, MD.

## DATA AVAILABILITY STATEMENT

Proteomics data have been deposited to the ProteomeXchange Consortium via the PRIDE partner repository [55] with the identifier PXD045193.

